# Extracellular adenosine induces hypersecretion of IL-17A by T-helper 17 cells through the adenosine A2a receptor to promote neutrophilic inflammation

**DOI:** 10.1101/2021.04.29.441713

**Authors:** Mieko Tokano, Sho Matsushita, Rie Takagi, Toshimasa Yamamoto, Masaaki Kawano

## Abstract

Extracellular adenosine, produced from ATP secreted by neuronal or immune cells, may play a role in endogenous regulation of inflammatory responses. However, the underlying molecular mechanisms are largely unknown. Here, we show that adenosine primes hypersecretion of interleukin (IL)-17A by CD4^+^ T cells via T cell receptor activation. This hypersecretion was also induced by an adenosine A2a receptor (A2aR) agonist, PSB0777. In addition, an A2aR antagonist (Istradefylline), and inhibitors of adenyl cyclase, protein kinase A, and cAMP response element binding protein (which are signaling molecules downstream of the Gs protein coupled with the A2aR), suppressed IL-17A production, suggesting that activation of A2aR induces IL-17A production by CD4^+^ T cells. Furthermore, immune subset studies revealed that adenosine induced hypersecretion of IL-17A by T-helper (Th)17 cells. These results indicate that adenosine is an endogenous modulator of neutrophilic inflammation. Administration of an A2aR antagonist to mice with experimental autoimmune encephalomyelitis led to marked amelioration of symptoms, suggesting that suppression of adenosine-mediated IL-17A production is an effective treatment for Th17-related autoimmune diseases.

## Introduction

T-helper (Th)17 cells are a subset of T-helper cells induced by stimulation of naïve CD4^+^ T cells with both tumor growth factor (TGF)-β and interleukin (IL)-6 in the presence of T cell receptor signaling. IL-17A production by Th17 cells increases neutrophilic inflammation (1–3); however, not all neutrophilic inflammatory diseases are explained by known Th17 responses (4). Indeed, it is likely that as-yet-unknown Th17 or neutrophilic inflammatory responses occur. Here, we show that adenosine induces IL-17A production by CD4^+^ T cells directly. Extracellular adenosine is one of the first “signals” identified during regulation of a large number of physiological and pathological processes, including bulging of an artery (5), sleep promotion (6), and regulation of nerve action (7, 8). Extracellular adenosine is produced from secreted ATP that undergoes rapid stepwise dephosphorylation by ectonucleotidases such as the E-NTPDase CD39, which converts ATP or ADP to ADP or AMP, respectively, and the 5’-nucleotidase CD73, which dephosphorylates AMP to adenosine (9); both CD39 and CD73 are expressed by activated CD4^+^ T cells and antigen presenting cells (APCs) (10, 11). Extracellular adenosine stimulates adenosine receptors (A1R, A2aR, A2bR, and A3R) belonging to a superfamily of membrane proteins called the G protein-coupled receptor family of class A seven-transmembrane domain receptors. A2aR and A2bR signal the Gs protein to trigger cAMP synthesis, which in turn activates adenyl cyclase (AC), protein kinase A (PKA), and cAMP response element binding protein (CREB). By contrast, A1R and A3R signal the Gi protein to trigger cAMP degradation. In addition, A2bR also signals the Gq protein, which in turn activates phospholipase C. In an immunological context, adenosine receptors are expressed by various immune cells, including T cells and APCs (12). It is also suggested that adenosine stimulates neutrophil chemotaxis and phagocytosis via A1R and A3R (13). In addition, adenosine induces Th17 differentiation by activating A2bR on CD4^+^ T cells (14). By contrast, several reports suggest that adenosine suppresses Th17 differentiation via activation of A2aR on CD4^+^ T cells (15–17). Considering that the G protein downstream of A2aR is Gs, and those of A2bR are Gs and Gq (5), it is assumed that induction of A2bR-mediated Th17 differentiation is induced through simultaneous activation of Gq and suppression of Gs, or via other unknown mechanisms. Therefore, the precise molecular mechanism(s) underlying the effect of adenosine on Th cells is unclear. Here, we show that adenosine promotes IL-17A production in a two-way mixed lymphocyte reaction (MLR). In addition, an A2aR agonist (PSB0777) induced IL-17A production, and an A2aR antagonist (Istradefylline) inhibited production, induced by adenosine, suggesting that activation of A2aR plays a role in adenosine-mediated IL-17A production. This notion was further supported by the observation that inhibitors of AC, PKA, and CREB, which are signaling molecules downstream of the Gs protein (18), also suppressed adenosine-mediated IL-17A production. Immune subset studies suggested that Th17 cells play a role in adenosine-mediated hypersecretion of IL-17A. Administration of an A2R antagonist to mice with experimental autoimmune encephalomyelitis (EAE) (19, 20) markedly ameliorated symptoms. Taken together, the data indicate that adenosine-dependent hypersecretion of IL-17A by Th17 cells contributes not only to antibacterial defense but also to neutrophilic autoimmune diseases, and that suppressing this process may be an effective therapy for the latter.

## Methods

### Mice

BALB/c mice were obtained from Japan SLC, Inc. SJL/J mice were obtained from Charles River Laboratories Japan, Inc. Mice were housed in appropriate animal care facilities at Saitama Medical University and handled according to international guidelines for experiments with animals. All experiments were approved by the Animal Research Committee of Saitama Medical University.

### Two-way MLR

Splenic lymphocytes were collected by lyzing tissue in a Dounce homogenizer, followed by layering over Ficoll Paque (GE health care, Chicago, IL, USA), as described previously (21). BALB/c splenic lymphocytes (3 × 10^6^) were mixed with SJL/J splenic lymphocytes (3 × 10^6^) in 2 mL of DMEM medium containing 10% FCS, 100 U/ml penicillin, 100 μg/mL streptomycin, 2 mM L-glutamine, 1 mM sodium pyruvate, and 50 μM 2-mercaptoethanol (D10 medium) in 12 well plates in the presence of adenosine (0–1 mM) (Sigma, St. Louis, MO, USA); in the presence of each adenosine receptor agonist (0–10 μM) (A1R: 2-Chloro-N6-cyclopentyladenosine (CCPA), Tocris, Bristol, UK; A2aR: PSB0777, Tocris; A2bR: BAY 60-6583, Tocris; A3R: HEMADO, Tocris); in the presence of A2aR antagonist (0–1 nM) (Istradefylline, Sigma) plus adenosine (100 μM); in the presence of an AC inhibitor (0–1 μM) (MDL-12330A, Enzo Life Sciences, Farmingdale, NY, USA), a PKA inhibitor (0–1 μM) (H-89, Tocris) plus adenosine (100 μM), or a CREB inhibitor (0.1–0.6 μM) (666-15, Cayman Chemical Company, Ann Arbor, MI, USA) plus adenosine (100 μM) (to inhibit A2aR signaling); in the presence of an activator of AC (0.06–0.6 μM) (Forskolin, Sigma); or in the presence of a CD39 inhibitor (0–1 μM) (ARL67156, Tocris) or a CD73 inhibitor (0–1 μM) (adenosine 5’-(α, β-methylene) diphosphate (AMP-CP; Tocris) plus ATP (100 μM) (GE healthcare). After mixing, the plates were incubated for 7 days at 37°C. The supernatants were collected for use in cytokine ELISAs.

### Isolation of CD4^+^, CD4^+^CD62L^+^ T cells, and B cells

CD4^+^ T cells within the BALB/c and SJL/J splenocyte populations were isolated from the mixture prepared previously after rupturing red blood cells (22). Cells were isolated by positive selection of CD4^+^ T cells using magnetic-activated cell sorting (MACS) (Miltenyi Biotec, Bergisch Gladbach, Germany), according to the manufacturer’s instructions. CD4^+^CD62L^+^ T cells were isolated from BALB/c splenocytes by a combination of negative and positive selection by MACS. During positive selection of CD62L^+^ T cells, negatively isolated CD4^+^ T cells were collected as the flow through fraction. B cells were isolated from BALB/c splenocytes by positive selection using MACS. Cells were resuspended in 1 mL of D10 medium and counted. The purity and viability of CD4^+^ T cells, CD4^+^CD62L^+^ T cells, and B cells were >90% (Sup. Fig. 1). The purity and viability of cells in the flow through fraction collected during isolation of CD4^+^CD62L^+^ T cells are shown in Supplementary Figure 1.

**Fig. 1.**
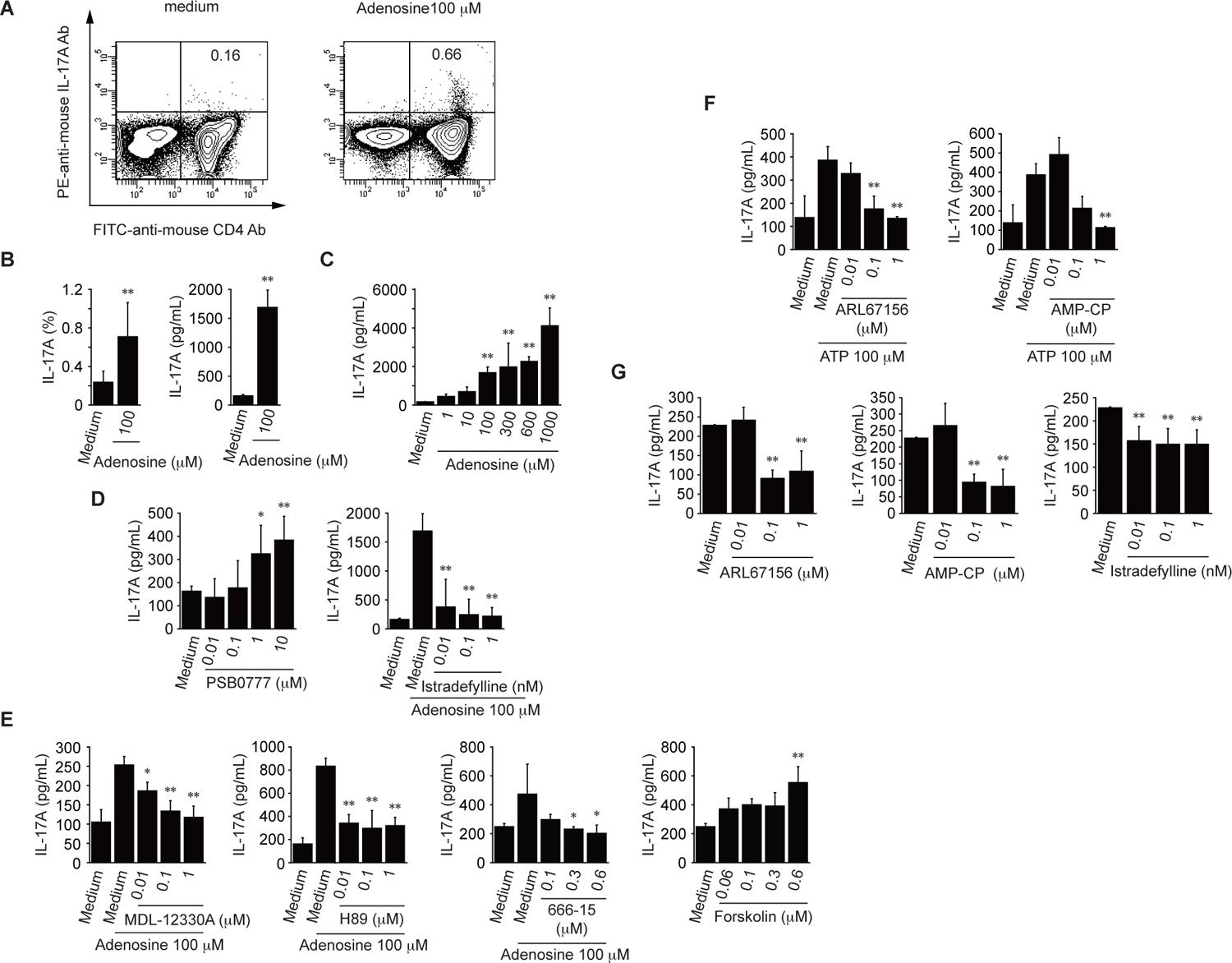
Adenosine induces hypersecretion of IL-17A by CD4^+^ T cells in an MLR. **A–C**, An MLR was performed for 7 days in the presence of adenosine (0–1 mM). After 7 days, cells were stained with anti-CD4 (x-axis) and IL-17A (y-axis) antibodies (Abs), followed by flow cytometry analysis (**A**, n (number of repeat experiments) = 4). The percentage of IL-17A-producing CD4^+^ T cells within the total CD4^+^ T cell population is shown (**B**, n = 6–9). Cells supernatants were analyzed in an IL-17A ELISA (**C**, n = 6–9). **D**, The effects of the A2aR on IL-17A production in the presence of PSB0777 (an A2aR agonist) (*left*, n = 6–9), Istradefylline (an A2aR antagonist) plus adenosine (100 μM) (*right*, n = 6–9). **E**, The effects of A2aR signaling on IL-17A production in the presence of MDL-12330A (an adenyl cyclase inhibitor) plus adenosine (100 μM) (*first panel*, n = 4–6), H-89 (a protein kinase A inhibitor) plus adenosine (100 μM) (*second panel*, n = 4–6), 666-15 (a cAMP response element binding protein inhibitor) plus adenosine (100 μM) (*third panel*, n = 4–6), and Forskolin (an activator of adenyl cyclase) (*fourth panel*, n = 4–6) were analyzed in an IL-17A ELISA. **F**, The effects of CD39/CD73 inhibitors on IL-17A production in the presence of ARL67156 (a CD39 inhibitor) plus ATP (100 μM) (*left*, n = 4 – 6) or AMP-CP (a CD73 inhibitor) plus ATP (100 μM) (*right*, n = 4–6) were analyzed in an IL-17A ELISA. **G**, The effects of an A2aR antagonist and CD39/CD73 inhibitors on basal IL-17A production in the presence of Istradefylline (*left*, n = 4– 7), ARL67156 (*center*, n = 4–7), or AMP-CP (*right*, n = 4–7) were analyzed in an IL-17A ELISA. Data are expressed as the mean ± standard deviation (SD) and were compared using an unpaired Student’s t-test (**B**) or one-way ANOVA with Tukey’s post-hoc test (**C**–**G**). *P < 0.05 and **P < 0.01, compared with medium (**C**, **D**, *left*; **E**, *fourth panel*; and **G**), adenosine (100 μM) (**D**, *right*, **E**, *first*–*third panels*), or ATP (100 μM) (**F**).

### Cell sorting

BALB/c CD4^+^ T cells (prepared as described above) were labeled for 30 min at 4°C with PE-conjugated anti-mouse CCR3, CCR5, CCD6, CD25, or CD62L antibodies (BioLegend). The cells were then washed and sorted using a FACS Aria II flow cytometer (BD Biosciences). The purity and viability of the sorted cells are Supplementary Figure 1.

### CD3/CD28 stimulation

CD4^+^ T cells (1 × 10^6^) were stimulated for 7 days at 37°C with anti-mouse CD3 (BioLegend; 1 μg/mL) and CD28 (BioLegend; 0.5 μg/mL) antibodies (CD3/CD28) in the presence of adenosine (0–1 mM), each adenosine receptor agonist (0–10 μM; A1R: 2-Chloro-N6-cyclopentyladenosine (CCPA), Tocris, Bristol, UK; A2aR: PSB0777, Tocris; A2bR: BAY 60-6583, Tocris; and A3R: HEMADO, Tocris), an A2aR antagonist (Istradefylline; 0–1 nM) plus adenosine (600 μM), an AC inhibitor (0–1 μM; MDL-12330A), a PKA inhibitor (0–1 μM) (H-89), a CREB inhibitor (0.1–0.6 μM; 666-15) plus adenosine (600 μM), or an activator of AC (0.06–0.6 μM; Forskolin) in 500 μL of D10 medium. After cell sorting, cells (3 × 10^5^) were plated in 24 well plates and stimulated for 7 days at 37°C with anti-mouse CD3 (1 μg/mL) and CD28 (0.5 μg/mL) antibodies in the presence of adenosine (600 μM) in 100 μL of D10 medium. After stimulation, the supernatants were collected for use in cytokine ELISAs.

### Adenosine or ATP ELISAs

MLR was performed by mixing BALB/c lymphocytes (6 × 10^6^) with SJL/J lymphocytes (6 × 10^6^) in a 15 mL tube for 0–24 h at 37°C in the presence of a CD39 inhibitor (ARL67156) (0–1 μM) and a CD73 inhibitor (AMP-CP) (0–1 μM) in 200 μL of D10 medium. CD3/CD28 stimulation was performed for 24 h at 37°C in a 15 mL tube by incubating BALB/c CD4^+^ T cells (1 × 10^7^) with anti-mouse CD3 (1 μg/mL) and CD28 (0.5 μg/mL) antibodies plus ARL67156 (1 μM) or AMP-CP (1 μM) in 200 μL of D10 medium. Simulation with LPS or an anti-mouse CD40 antibody (Ab) was performed for 24 h at 37°C in a 15 mL tube; briefly, BALB/c B cells (1 × 10^7^) or BALB/c bone marrow (BM)-derived dendritic cells (BM-DCs) (1 × 10^7^), generated from mouse bone marrow cells as described previously (23), were incubated with LPS (Sigma; 0.5 μg/mL) or anti-mouse CD40 Ab (5 μg/mL; BioLegend) plus ARL67156 (1 μM) or AMP-CP (1 μM) in 200 μL of D10 medium. The purity and viability of BM-DCs were >90% (Sup. Fig. 1). After incubation, the supernatants were collected and tested in adenosine or ATP ELISAs (Biovision, Milpitas, CA, USA).

### Differentiation of naïve CD4^+^ T cells

Naïve CD4^+^ T cells (3 × 10^5^) in 500 μL of D10 medium in 24 well plates were stimulated for 7 days at 37°C with anti-mouse CD3 (1 µg/mL) and CD28 (0.5 mg/mL) antibodies plus mouse IL-6 (20 ng/mL) (Peprotech, Rocky Hill, NJ), and human TGF-β1 (2 ng/mL) (Peprotech) in the presence of an A2aR antagonist (Istradefylline) (0–1 nM). After incubation, the supernatants were collected for use in cytokine ELISAs, and total RNA was extracted from the cells using a Sepasol-RNA I Super G (Nacalai Tesque, Kyoto, Japan) for quantitative PCR. In another experiment, cells in 500 μL of D10 medium were stimulated for another 7 days at 37°C with anti-mouse CD3 (1 µg/mL) and CD28 (0.5 μg/mL) antibodies in the presence of Istradefylline (0.1–1 nM) plus adenosine (600 μM). Supernatants were collected for use in cytokine ELISAs.

### Quantitative PCR

cDNA was synthesized from total RNA prepared using Superscript IV VILO master mix (Invitrogen, Waltham, MA, USA), according to the manufacturer’s instructions. Quantitative PCR (qPCR) was performed as described previously (24) using the primer pairs described below, Power SYBR green PCR Master mix (Applied Biosystems, Waltham, MA, USA), and a QuantStudio 12K Flex (Applied Biosystems) equipped with QuantStudio 12K Flex software (Applied Biosystems). Then, the ⊗Ct was calculated. The following forward and reverse primers, respectively, were used (all from Eurofins Scientific, Luxembourg): GAPDH, 5’-AGCTTGTCATCAACGGGAAG-3’ and 5’-TTTGATGTTAGTGGGGTCTCG-3’; IL-17A, 5’-TGTGAAGGTCAACCTCAAAGTCT-3’ and 5’-GAGGGATATCTATCAGGGTCTTCAT-3’.

### EAE model

EAE was induced as described previously (19). Briefly, SJL/J mice received a subcutaneous inguinal injection (100 μg/mouse) of the proteolipid protein (PLP) peptide (PLP139-151, Tocris) emulsified in complete Freund’s adjuvant (CFA) containing mycobacterium tuberculosis H37Ra (100 μg/mouse; Difco, Detroit, MI, USA). Mice also received oral water or an A2aR antagonist (Istradefylline; 6 μg/mouse) once every 2 days from Day –7 to Day +18 after immunization with PLP peptide (Day 0). Mice were examined daily for signs of EAE, which were graded as described (25).

### Histological analysis of spinal cord samples

On Day 18 post-induction, histological analysis was performed by excising the lumbar, thoracal, and cervical parts of the spinal cord. Paraffin-embedded spinal cord sections were stained with hematoxylin and eosin. T cells were labeled with an anti-mouse CD3ε Ab (Cell Signaling technology, Danvers, MA, USA). Tissue sections were observed using the light microscopy mode of an inverted fluorescence phase-contrast microscope (BZ-9000, Keyence, Osaka, Japan).

### Peptide pulse assay

At 7 days post-subcutaneous immunization with PLP peptide emulsified in CFA, splenocytes (2 × 10^6^ in 200 μL of D10 medium) were seeded in 96-well round plates and pulsed for 3 days at 37°C with PLP peptide (10 μM) in the presence of adenosine (600 μM) and an A2aR antagonist (Istradefylline; 0–1 nM). For the oral administration experiments using the A2a antagonist, splenocytes (2 × 10^6^ in 200 μL of D10 medium) or inguinal lymph node lymphocytes, prepared as described previously (26) (1 × 10^6^ in 200 μL of D10 medium) were seeded in 96-well round plates and pulsed for 3 days at 37°C; this was done at 7 days post-subcutaneous immunization with PLP peptide emulsified in CFA. Mice also received oral water, an A2aR agonist (PSB0777) (6 μg/mouse), or an A2aR antagonist (Istradefylline; 6 μg/mouse) once every 2 days from Day – 7 to Day +7 after immunization with PLP peptide (Day 0). At 18 days post-subcutaneous immunization with PLP peptide emulsified in CFA, spinal cord cells prepared as described previously (26) (100 μL of resuspended in 500 μL of D10 medium/mouse) were seeded in 96-well round plates and pulsed for 3 days at 37°C with PLP peptide (10 μM). The supernatants were collected for use in cytokine ELISAs.

### Flow cytometry analysis

The MLR was performed for 7 days in the presence or absence of adenosine (100 μM) (as described above). For the CD4^+^IL-17A^+^ study, cells were blocked with anti-mouse CD16/CD32 antibodies (BioLegend, San Diego, CA, USA) and stained for 30 min at 4°C with a fluorescein isothiocyanate (FITC)-conjugated anti-mouse CD4 Ab (BioLegend). The cells were then fixed, permeabilized, and stained with a phycoerythrin (PE)-conjugated anti-mouse IL-17A Ab (BioLegend). For the Th subset study, after 0, 3, and 7 days of the MLR assay, cells were blocked with anti-mouse CD16/CD32 antibodies (BioLegend) and then stained for 30 min at 4°C with a FITC-conjugated anti-mouse CD4 Ab (BioLegend) and PE-conjugated anti-mouse CCR5 (a Th1 marker (27)), CCR3 (a Th2 marker (28)), or CCR6 (a Th17 marker (29, 30)) antibodies (BioLegend). For the MHC, costimulatory molecule, and A2aR expression studies in APCs, BALB/c B cells (3 × 10^5^) or BALB/c BM-DCs (3 × 10^5^) seeded in the wells of a 96-well in 200 μL of D10 medium were incubated with LPS (0.5 μg/mL; Sigma) or an anti-mouse CD40 Ab (5 μg/mL) (BioLegend) for 1 day. Next, supernatants were collected for cytokine ELISAs, or blocked with anti-mouse CD16/CD32 antibodies (BioLegend) and then stained for 30 min at 4°C with FITC-conjugated anti-mouse CD80, CD86, or CD40 Abs and corresponding isotype controls (BioLegend) (for the costimulatory study), with PE-conjugated anti-mouse H-2 or I-A/I-E Ab and corresponding isotype controls (BioLegend) (for the MHC study), or fixed, permeabilized, and stained with a PE-conjugated anti-A2aR Ab (Novus Biologicals, Minneapolis, MN, USA) and corresponding isotype control (for the A2aR expression study). For the A2aR expression study in CD4^+^ T cells, CD4^+^ T cells (1 × 10^6^) were stimulated for 7 days at 37°C with anti-mouse CD3 (1 µg/mL) and CD28 (0.5 μg/mL) antibodies in the presence of adenosine (0–1 mM). At 0–7 days post-stimulation, cells were blocked with anti-mouse CD16/CD32 antibodies (BioLegend) and then stained for 30 min at 4°C with a FITC-conjugated anti-mouse CD4 Ab (BioLegend) before fixation, permeabilization, and staining with a PE-conjugated anti-A2aR Ab (Novus Biologicals). For EAE analysis at 18 days post-subcutaneous immunization with PLP peptide emulsified in CFA, splenocytes (1 × 10^6^), inguinal lymph node lymphocytes (1 × 10^6^), and spinal cord cells (100 μL of resuspended in 500 μL of D10 medium/mouse) were blocked with anti-mouse CD16/CD32 antibodies (BioLegend) and then stained for 30 min at 4°C with a FITC-conjugated anti-mouse CD4 Ab (BioLegend) and PE-conjugated anti-mouse CCR5 (a Th1 marker (27)), CCR3 (a Th2 marker (28)), or CCR6 (a Th17 marker (29, 30)) antibodies (BioLegend). Finally, the cells were washed and analyzed on a FACSCanto II flow cytometer (BD Biosciences, Franklin Lakes, NJ) using FACSDiva acquisition software (BD Biosciences).

### Cytokine ELISAs

The concentrations of IFN-γ, IL-5, IL-6, IL-17A, IL-17F, IL-22, IL-23, and TGF-β1, in cell supernatants were measured using specific ELISA kits (DuoSet Kit, R&D, Minneapolis, MN, USA). Any value below the lower limit of detection (15.6 pg/mL) was set to 0. No cytokine cross-reactivity was observed within the detection ranges of the kits. If necessary, samples were diluted appropriately so that the measurements fell within the appropriate detection range for each cytokine.

### Statistical analysis

Differences between two groups were analyzed using an unpaired Student’s t-tests. Differences between three or more groups were analyzed using one-way ANOVA with Tukey’s post-hoc test. Clinical scores were analyzed using a non-parametric Mann-Whitney *U*-test. All calculations were performed using KaleidaGraph software (Synergy software, Reading, PA, USA). A P value < 0.05 was considered statistically significant.

## Results

### Adenosine promotes IL-17A production by CD4^+^ T cells in an MLR

First, we analyzed the effect of adenosine on CD4^+^ T cells during T cell-APC interactions in an MLR (31). We found that CD4^+^ T cells exposed to adenosine secreted IL-17A in a dose-dependent manner (Fig. 1A–C). Since both agonist-mediated IL-17A production and antagonist-mediated suppression of adenosine-mediated IL-17A production were observed in the presence of an adenosine A2aR agonist (PSB0777) and an antagonist (Istradefylline), respectively (Fig. 1D and Sup. Fig. 2A), we hypothesized that IL-17A in the MLR was produced by CD4^+^ T cells stimulated via the A2aR. This notion was supported by the finding that AC, PKA, and CREB inhibitors of signaling molecules downstream of the A2aR also suppressed IL-17A production (Fig. 1E). Consistent with this notion, activation of A2aR downstream of AC upregulated IL-17A production (Fig. 1E). In the MLR, the Th1 and Th17 cell populations increased slightly but significantly (by approximately 1.7-fold) in the presence of 100 µM adenosine at Day 7 (Sup. Fig. 2C). Considering that 100 µM adenosine increased IL-17A secretion by approximately 10-fold compared with medium (Fig. 1B), it appears that adenosine-mediated upregulation of IL-17A is the main contributor to upregulation of the IL-17A production by CD4^+^ T cells. Furthermore, ATP induced IL-17A production in the MLR, which was suppressed by the A2aR antagonist and by inhibitors of CD39 and CD73 (Fig. 1F), suggesting that adenosine plays a role in IL-17A production. Since the A2aR antagonist and inhibitors of CD39 and CD73 also inhibited production of IL-17A in the MLR (Fig. 1G), we postulate that *de novo* adenosine production is induced in the MLR.

**Fig. 2.**
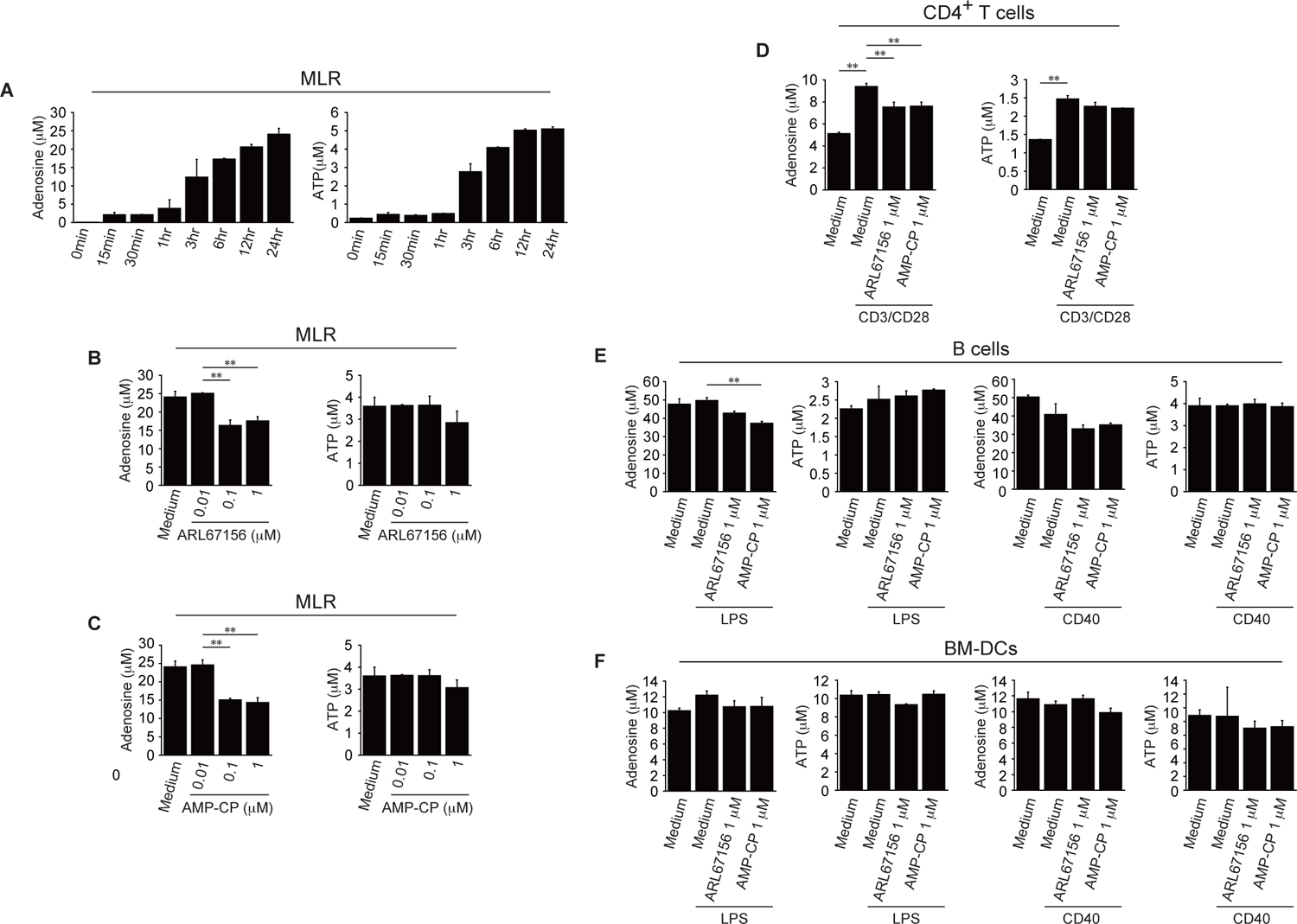
Adenosine production in the MLR. **A**, Concentrations of adenosine or ATP in MLR supernatants were measured in an adenosine (*left*, n = 4–6) or ATP (*right*, n = 4–6) ELISA (0–24 h). **B** and **C**, The effects of CD39 (**B**, ARL67156, n = 4–6) and CD73 (**C**, AMP-CP, n = 4–6) inhibitors on production of adenosine or ATP in the MLR were analyzed in an adenosine (*left*) or ATP (*right*) ELISA. **D, E** and **F**, Levels of adenosine and ATP in the supernatants of CD3/CD28-stimulated CD4^+^ T cells (**D**, n = 4), LPS- or anti-mouse CD40 Ab-stimulated B cells (**E**, n = 4), and LPS- or anti-mouse CD40 Ab-stimulated BM-DCs (**F**, n = 4) were analyzed in adenosine and ATP ELISAs at 24 h post-stimulation in the presence of CD39 or CD73 inhibitors. Data are expressed as the mean ± SD and were compared using one-way ANOVA with Tukey’s post-hoc test. *P < 0.05 and **P < 0.01, compared with medium (**B** and **C**), CD3/CD28 stimulation (**D**), or LPS or anti-mouse CD40 Ab stimulation (**E** and **F**).

### Adenosine production in the MLR

CD39 and CD73 expressed on the surface of endothelial cells (32, 33) and immune cells (10, 11), including T cells and DCs, are critical for production of adenosine from ATP. Since we found that inhibitors of CD39 and CD73 inhibited basal IL-17A production in the MLR (Fig. 1G), we next addressed the source of adenosine production in the MLR. As shown in Figure 2A, production of adenosine and ATP was time-dependent. In accordance with suppression of IL-17A production, inhibitors of CD39 and CD73 suppressed adenosine production at 24 h after the start of the MLR (Fig. 2B and C). Since the MLR induces activation of CD4^+^ T cells by APCs (31), we also addressed adenosine production by activated CD4^+^ T cells, B cells, and BM-DCs (Fig. 2D–F). Production of both adenosine and ATP by activated CD4^+^ T cells, B cells, and BM-DCs was observed; inhibitors of CD39 and CD73 suppressed production by activated CD4^+^ T cells although production of both adenosine and ATP by CD4^+^ T cells, B cells, and BM-DCs was observed without LPS or anti CD40 Ab stimulation (Fig. 2D). This indicates the possibility that adenosine stimulation is mediated by the CD4^+^ T cell to other immune cell contact and adenosine produced by CD4^+^ T cells or other immune cells such as B cells may induce adenosine-mediated IL-17A hypersecretion by CD4^+^ T cells in the MLR.

### Th17 cells hypersecrete IL-17A in the presence of anti-CD3/CD28 antibodies and adenosine

Next, we tried to identify the Th subset that generated IL-17A in the presence of adenosine. First, we confirmed that CD4^+^ T cells expressed the A2aR and secreted IL-17A by stimulating them with agonistic anti-CD3/CD28 antibodies in the presence of adenosine (Fig. 3A). Time course studies showed that adenosine-mediated IL-17A production was detected from 3 days post-CD3/CD28 stimulation (Fig. 3B and Sup. Fig. 3). Administration of adenosine within 6 h of Ab stimulation triggered IL-17A production; however, administration at 24 h post-stimulation did not (Fig. 3C). We noticed that A2aR expression by CD4^+^ T cells was upregulated significantly (2–4.7 fold) by anti-mouse CD3 and CD28 Abs at 1–7 days after stimulation, and that this upregulation was further (slightly but significantly) enhanced (by approximately 1.5-fold) by increasing amounts of adenosine (Sup. Fig. 4A). As in the MLR, CD4^+^ T cells also produced IL-17A after stimulation with anti-CD3/CD28 antibodies in the presence of an A2aR agonist (Fig. 3D and Sup. Fig. 2B). Adenosine-mediated IL-17A production by CD3/CD28-stimulated CD4^+^ T cells was suppressed by an A2aR antagonist (Fig. 3D and Sup. Fig. 2B). This was supported by data showing that inhibitors of signaling molecules downstream of the A2R also suppressed IL-17A production, whereas activation of A2aR downstream of AC upregulated the production (Fig. 3E). Production of other Th17-related cytokines was also induced by adenosine through the A2aR (Fig. 4). This again suggests that activated CD4^+^ T cells produce IL-17A upon A2aR activation. Furthermore, we noticed that CD4^+^CD62L^-^, but not CD4^+^CD62L^+^, cells produced IL-17A after CD3/CD28 stimulation in the presence of adenosine, suggesting that adenosine induces IL-17A production by effector Th cells (Fig. 3F). Therefore, we performed immune subset studies after separating CD4^+^ T cells using anti-chemokine receptor (CCR) antibodies. As shown in Figure 3G, CD4^+^CCR6^hi^ T cells produced IL-17A upon stimulation of CD3/CD28, and production was strongly upregulated in the presence of adenosine. Since CCR6 is a typical marker of Th17 cells (29, 30), this suggests that activated Th17 cells hypersecrete IL-17A in the presence of adenosine.

**Fig. 3.**
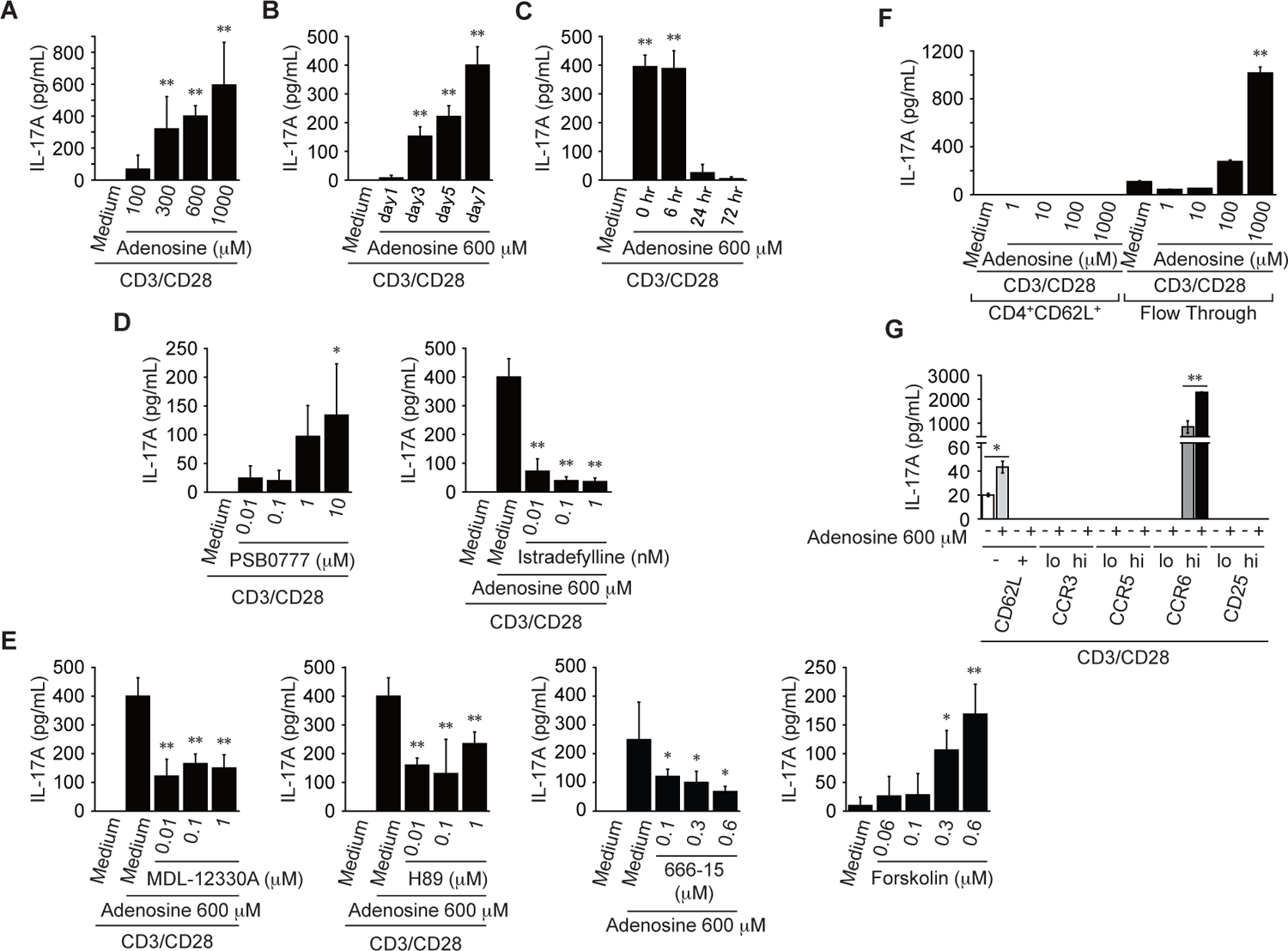
Adenosine induces hypersecretion of IL-17A by Th17 cells. **A** and **B**, CD4^+^ T cells were stimulated for 1–7 days with anti-CD3/CD28 antibodies in the presence of adenosine (0–1 mM). After stimulation, the supernatants were analyzed in an IL-17A ELISA (n = 4–6). **C**, Adenosine (600 μM) was added at 0–3 days after CD3/CD28 stimulation. At 7 days post-CD3/CD28 stimulation, supernatants were analyzed in an IL-17A ELISA (n = 4). **D**, Effects of the A2aR on IL-17A production in the presence of PSB0777 (an A2aR agonist) (*left*, n = 4–6), Istradefylline (an A2aR antagonist) plus adenosine (600 μM) (*right*, n = 4–6). **E**, Effects of A2aR signaling on IL-17A production in the presence of MDL-12330A (an adenyl cyclase inhibitor) plus adenosine (600 μM) (*first panel*, n = 4–6), or H-89 (a protein kinase A inhibitor) plus adenosine (600 μM) (*second panel*, n = 4–6), 666-15 (a cAMP response element binding protein inhibitor) plus adenosine (600 μM) (*third panel*, n = 4–6), or Forskolin (an activator of adenyl cyclase) (*fourth panel*, n = 4–6) were analyzed in an IL-17A ELISA. **F**, CD4^+^CD62L^+^ and CD4^+^CD62L^+^FT cells were stimulated for 7 days with anti-CD3/CD28 antibodies in the presence of adenosine (0–1 mM). After 7 days, supernatants were analyzed in an IL-17A ELISA (n = 4). **G**, Subsets of CD4^+^ T cells were stimulated for 7 days by anti-CD3/CD28 antibodies in the presence of adenosine (600 μM) after isolation of each CCR cell type (high (hi) and low (lo) expression) (n = 4). Data are expressed as the mean ± SD and were compared using an unpaired Student’s t-test (**G**) or one-way ANOVA with Tukey’s post-hoc test (**A–F**). *P < 0.05 and **P < 0.01, compared with CD3/CD28 stimulation (**A**–**C**, **D**, *left;* **E***, fourth panel;* and **F**) or CD3/28 stimulation plus adenosine (600 μM) (**D**, *right* and **E***, first*–*third panels*).

**Fig. 4.**
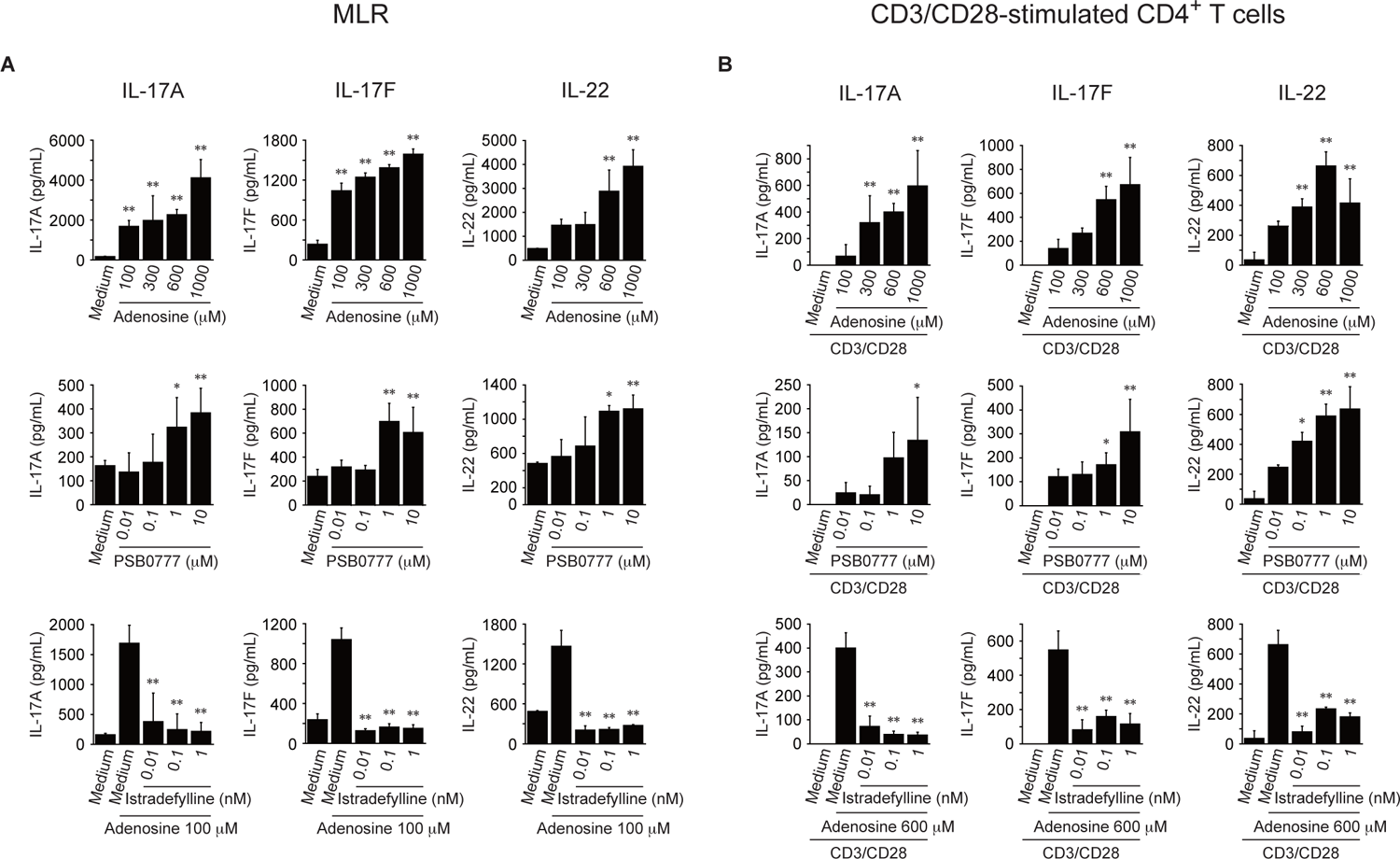
Effect of adenosine and an A2aR antagonist on production of Th17-related cytokines. **A,** An MLR was performed for 7 days in the presence of adenosine (0–1 mM) (*top row*), PSB0777 (an A2aR agonist) (*second row*), or Istradefylline (an A2aR antagonist) plus adenosine (100 μM) (*bottom row*). At 7 days post-incubation, the supernatants were analyzed in an IL-17A (*left*), IL-17F (*center*), or IL-22 (*right*) ELISA (n = 6–9). **B,** CD4^+^ T cells were stimulated with an anti-CD3/CD28 antibody for 7 days in the presence of adenosine (600 μM) (*top row*), PSB0777 (an A2aR agonist) (*second row*), or Istradefylline (an A2aR antagonist) plus adenosine (600 μM) *(bottom row*). After stimulation, supernatants were analyzed in IL-17A (*left*), IL-17F (*center*), and IL-22 (*right*) ELISAs (n = 4–6). Data are expressed as the mean ± SD and were compared using one-way ANOVA with Tukey’s post-hoc test. *P < 0.05 and **P < 0.01, compared with medium or CD3/CD28 stimulation (*top and second row of panels*) or adenosine (100 μM) or CD3/CD28 stimulation plus adenosine (600 μM) (*third row of panels*).

To address the effect of adenosine on APCs, we analyzed production of IL-6, IL-23, and, TGF-β1 by B cells and BM-DCs (Sup. Fig. 4B). Also, we analyzed the effect of adenosine on expression of CD80/CD86/CD40, MHC class I/class II, and A2aR by B cells and BM-DCs (Sup. Fig. 4C). Increasing amounts of adenosine suppressed IL-6 production by LPS-stimulated B cells. and IL-6 and IL-23 production by LPS-stimulated BM-DCs. Expression of CD80/CD86/CD40, MHC class I/class II, and A2aR by LPS or anti-CD40 Ab-stimulated-B cells and -BM-DCs in the presence of 100 µM and 600 µM adenosine was 0.70- and 1.31-fold higher than that by cells stimulated by LPS or anti-CD40 Ab alone. These data indicate that adenosine mainly affects CD4^+^ T cells to activate Th17 function.

### An adenosine A2aR antagonist ameliorates IL-17A-related autoimmune EAE responses

The above results raise the possibility that adenosine-mediated hypersecretion of IL-17A by Th17 cells contributes to Th17-related autoimmune diseases. This hypothesis is supported by a report showing that CD73 knockout mice are resistant to EAE (34), a Th17-mediated autoimmune disease (20). We expected, therefore, that A2aR antagonist-mediated suppression of Th17 responses should improve EAE. To address this, we examined the efficacy of an A2aR antagonist in EAE model SJL/J mice (19). EAE was induced by immunization of mice with an I-A^s^ restricted helper peptide derived from a myelin PLP peptide comprising amino acids 139– 151 (HSLGKWLGHPDKF). The peptide was emulsified in CFA. First, we confirmed that the A2aR antagonist suppressed adenosine-mediated IL-17A production by CD3/CD28-stimulated CD4^+^ T cells from SJL/J strain mice (Fig. 5A). The A2aR antagonist also significantly suppressed adenosine-mediated IL-17A production after differentiation of Th17 cells from naïve CD4^+^ T cells (Fig. 5B, *right*). The A2aR antagonist did not suppress IL-17A expression or production during differentiation of Th17 cells from naïve CD4^+^ T cells (Fig. 5B, *left and center*). By contrast, and in agreement with Figure 3G, adenosine administration did not induce IL-17A production during and after differentiation of Th1, Th2, and Treg cells from naïve CD4^+^ T cells (data not shown). Next, we pulsed splenocytes with the PLP peptide after immunization to confirm that IL-17A production was induced in a peptide-dependent manner, and that production was upregulated by adenosine. As expected, IL-17A production occurred in a peptide-dependent manner and was upregulated by adenosine (Fig. 5C). Furthermore, the A2aR antagonist suppressed production of IL-17A, suggesting that the A2aR antagonist inhibits IL-17A production by CD4^+^ T cells induced by immunization with the PLP peptide. We next addressed the effect of an oral A2aR antagonist on production of IL-17A by CD4^+^ T cells induced by immunization with the PLP peptide (Fig. 5D). Oral administration of PSB0777 or Istradefylline did not significantly induce or suppress IL-17A production by PLP peptide-pulsed splenocytes and inguinal lymph node lymphocytes. These results indicate that the A2aR antagonist may not inhibit PLP peptide-specific Th17-generation but rather suppress adenosine-mediated IL-17A hypersecretion by PLP peptide-specific Th17 cells. Since oral administration of PSB0777 did not up-regulate IL-17A production, adenosine-mediated IL-17A production may be saturated by *de novo* extracellular adenosine in the body. Hence, we expect that the A2aR antagonist may suppress adenosine-mediated IL-17A hypersecretion by PLP peptide-specific Th17 cells during migration to the inflammatory site, which may suppress EAE symptoms. Finally, the A2aR antagonist was administered orally to mice before and during EAE induction (Fig. 5E and F). As shown, the clinical scores of mice receiving the A2aR antagonist were markedly lower than those of control mice (receiving water) at 18 days post-immunization with the PLP peptide (Fig. 5E). Accordingly, histological studies showed that the numbers of central nervous system-infiltrating CD3^+^ cells in mice receiving the A2aR antagonist were much lower than those in mice receiving water (Fig. 5F). Consistent with these data, flow cytometry analysis showed that the proportion of CD4^+^ cells in total counts in the spinal cord were reduced significantly by the oral A2aR antagonist; in particular, the percentage of Th17 cells in the spinal cord was reduced significantly (Fig. 5G), suggesting the A2aR antagonist suppressed migration of CD4^+^ T cells to, and retained Th17 cells within, the spinal cord during EAE induction. Accordingly, PLP peptide-dependent production of IL-17A was observed in spinal cord cells after oral administration of water; production was suppressed significantly by the oral A2aR antagonist (Fig. 5H). These results suggest that the A2aR antagonist suppresses hypersecretion of IL-17A by PLP peptide-specific Th17-cells, resulting in impaired migration from the draining lymph nodes to the inflammatory site, and subsequent retention at the inflammatory site. These phenomena may suppress IL-17A-mediated EAE symptoms.

**Fig. 5.**
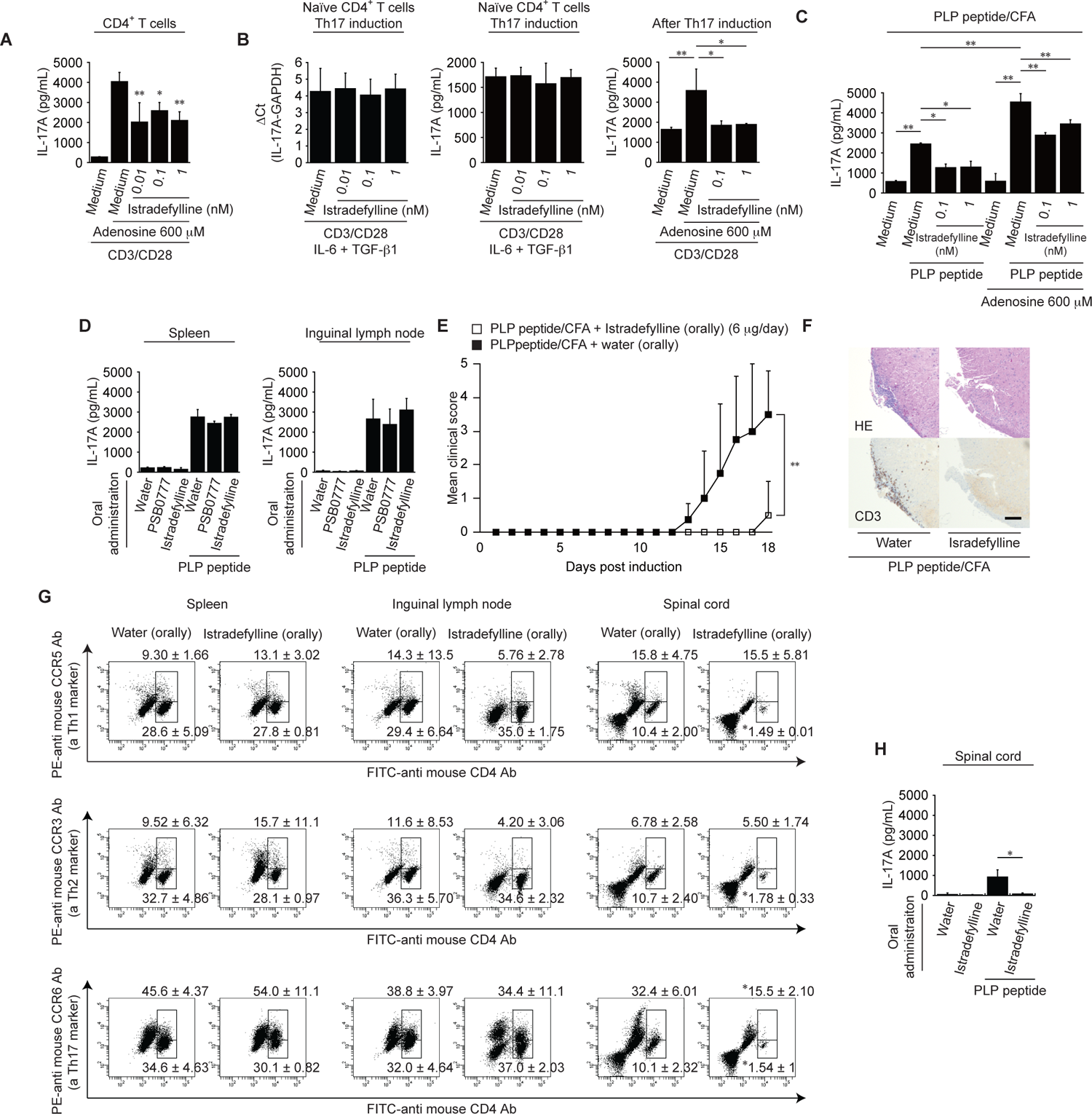
Suppression of adenosine-mediated hypersecretion of IL-17A ameliorates EAE. **A** and **B**, SJL/J CD4^+^ T cells were stimulated for 7 days by anti-CD3/CD28 antibodies and an A2aR antagonist (**A**, n = 4). Naïve CD4^+^ T cells were stimulated for 7 days by anti-CD3/CD28 antibodies in the presence of IL-6, TGF-β1, and Istradefylline (an A2aR antagonist) (0–1 nM) (**B**, *left*, n = 4). Alternatively, naïve CD4^+^ T cells were stimulated for 7 days by anti-CD3/CD28 antibodies in the presence of IL-6 and TGF-β1, followed by another 7 day incubation with anti-CD3/CD28 antibodies and Istradefylline (0–1 nM) (**B**, *right*, n = 4). After the stimulation, supernatants were analyzed in an IL-17A ELISA. **C and D,** Mice were immunized subcutaneously with PLP peptide emulsified in CFA (PLP peptide/CFA). Mice received oral water, an A2aR agonist (PSB0777; 6 μg/mouse), or an A2aR antagonist (Istradefylline; 6 μg/mouse) once every 2 days from Day –7 to Day +7 after immunization with PLP peptide/CFA (Day 0) (**D**). At 7 days post-immunization, splenocytes were incubated for 3 days with PLP peptide in the presence of Istradefylline and adenosine (600 μM) (**C**, n = 4) or in the presence or absence of PLP peptide (**D**, n = 4). After the incubation, supernatants were analyzed in an IL-17A ELISA. **E,** To induce EAE, SJL/J mice were immunized with PLP peptide/CFA. Before and post-immunization with the PLP peptide (Days 0 to 18), mice received oral Istradefylline (6 μg/mouse) or water once every 2 days. Clinical scores were recorded every day during EAE induction (**E**, n = 15). Data were obtained from five independent experiments (n = 3 mice/group). **F,** At 18 days post-immunization, spinal cord sections from mice administered oral Istradefylline (*right*) or water (*left*) were stained with hematoxylin and eosin (*upper panels*) or with an anti-mouse CD3 antibody (*lower panels*) (Scale bar, 100 μm) (**F**, n = 5). **G and H,** At 18 days post-immunization, splenocytes (spleen, *left*), inguinal lymph node lymphocytes (inguinal lymph node, *center*), and spinal cord cells (spinal cord, *right*) from mice administered oral water (*left* of each panel) or Istradefylline (*right* of each panel) were stained with anti-CD4 (x-axis) and CCR5 (a Th1 marker; top panels, y-axis), CCR3 (a Th2 marker; second panels, y-axis), or CCR6 (a Th17 marker; bottom panels, y-axis) antibodies (Abs), followed by flow cytometry analysis (**G**, n = 5). Values shown in the upper and lower right of each panel denote the percentage of CCR-positive counts within the total CD4-positive counts, and the percentage of CD4-positive counts within the total count, respectively. *P < 0.05. Also, spinal cord cells were incubated for 3 days in the presence or absence of PLP peptide (**H**, n = 4). After the incubation, supernatants were analyzed in an IL-17A ELISA. Data are expressed as the mean ± SD. Data were compared using one-way ANOVA with Tukey’s post-hoc test (**A**–**D, G, and H**), or using a non-parametric Mann-Whitney *U*-test (**E**). *P < 0.05 and **P < 0.01, compared with CD3/CD28 stimulation plus adenosine (600 μM) (**A**), CD3/CD28 stimulation plus IL-6 and TGF-β1 (**B**), PLP peptide pulse (**C**), oral water (**G**), or PLP peptide pulse (oral water) (**H**).

## Discussion

Here, we showed that adenosine induces hypersecretion of IL-17A by Th17 cells. Addition of adenosine (1 mM) to a two-way MLR increased IL-17A production to > 25 times the basal level; however, both basal production and increased production of IL-17A were suppressed by an A2aR antagonist, and by CD39/CD73 inhibitors. This indicates that hypersecretion of IL-17A in the presence of adenosine occurs by other mechanisms in addition to T cell-APC interactions. Since endothelial cells and nervous system cells also express CD39/CD73 (32,33,35,36) and produce adenosine (8, 37), and activated CD4^+^ T cells in the present study hypersecreted IL-17A at 6 h post-CD3/CD28 stimulation, it is possible that activated Th17 cells also receive adenosine from endothelial and neuronal cells to induce hypersecretion of IL-17A.

We hypothesize that adenosine affects multiple immunological events as follows: After APCs take up antigen they move to the secondary lymphoid organs. Followed by an innate immune response in the secondary lymphoid organs, APCs induce Th differentiation from naïve CD4^+^ T cells. In particular, APCs induce Th17 differentiation by secreting IL-6 and TGF-β1. Then, the transcription factor RORγt switches on the IL-17A gene by binding to the IL-17A promoter (38). During this response, adenosine rarely stimulates naïve CD4^+^ T cells due to low levels of A2aR expression (Sup. Fig. 4A). Following an adaptive immune response, APCs in the secondary lymphoid organs stimulates effector Th17 cells and drives expression of the IL-17A gene by recruiting transcription factors such as NF-κB and NFAT downstream of the TCR signaling pathway (39). In this response, A2aR expression is also upregulated (Sup. Fig. 4A), thereby increasing and sensitivity of A2aR signaling in response to adenosine. After stimulation, Th17 cells migrate toward the inflammatory site. During migration, activated Th17 cells receive adenosine from immune cells, neural cells, endothelial cells, or in an autocrine manner. During this process, Th17 cells further up-regulate expression of the IL-17A gene by recruiting transcription factors such as CREB downstream of the A2aR-cAMP-PKA pathway (40). Then, Th17 cells induce IL-17A hypersecretion at the inflammatory site. In humans, the transcription factor cAMP-responsive element modulator α increases transcription of the human *Il-17* gene by binding to the promoter region (41). The A2aR antagonist may suppress adenosine-mediated IL-17A hypersecretion by adenosine-stimulated Th17 cells at the inflammatory site and subsequently suppress neutrophil-mediated inflammatory responses and Th cell migration. This hypothesis is consistent with a previous EAE study showing that CD73 expression and adenosine receptor signaling are required for efficient entry of lymphocytes into the central nervous system during EAE development (34); our own data suggest that CD73 expression and adenosine receptor signaling may be required to induce IL-17A hypersecretion by Th17 cells around the inflammatory site. γδ T cells, which are categorized mainly into two subpopulations (CD8αα^+^ and CD8^−^ cells), express RORγt and are also a source of IL-17A (42, 43). We hypothesize that the A2aR antagonist will also suppress adenosine-mediated IL-17A production by adenosine-stimulated γδ T cells through A2aR signaling (44).

During the sequence of immunological events described above, TCR signaling up-regulates phosphodiesterase activity and suppresses cAMP signaling during the adaptive responses in the secondary lymphoid organs (45). The cAMP–PKA signaling pathway plays a major role in regulating immune responses, and cAMP is the most potent and acute inhibitor of T cell activation (45). Our data indicate that up-regulation of A2aR expression is induced after TCR stimulation (Sup. Fig. 4A); therefore, adenosine-mediated IL-17A hypersecretion is induced after stimulation of TCR signaling as seen in Fig. 3C. Therefore, extracellular adenosine may not block TCR signaling in the body. Also, a specific amount of Forskolin may not block TCR signaling, leading to subsequent upregulation of IL-17A secretion by TCR-activated CD4^+^ T cells (Fig. 1E and 3E). We hypothesize that extracellular adenosine-mediated IL-17A hypersecretion is induced after activation of TCR signaling during migration from the secondary lymphoid organs to the inflammatory site. In the MLR and the PLP peptide pulse experiment, both activated Th17 cells and adenosine-stimulated Th17 cells might produce IL-17A during the T-APC interaction. Thus, inhibitors of CD39/CD73 and an A2aR antagonist may suppress adenosine-mediated hypersecretion by adenosine-stimulated activated Th17 cells.

It is suggested that physiological concentrations of adenosine are lower than 1 μM; however, they can be increased by stimuli such as high K^+^ levels, electrical stimulation, glutamate receptor agonists, hypoxia, hypoglycemia, and ischemia (46). To obtain sufficient adenosine (> 100 μM) to trigger hypersecretion of IL-17A, activated Th17 cells may need to make contact with non-immune cells such as adenosine-producing endothelial cells (47) and neuronal cells (48) to form a microenvironment with a high adenosine concentration (as observed during T cell-APC interactions at immunological synapses) (49). Thus, A2aR antagonists, rather than CD39/CD73 inhibitors, might be more effective at inhibiting *de novo* adenosine-mediated hypersecretion of IL-17 by Th17 cells. A previous study suggests that intracellular adenosine is transported out of cells by efficient equilibrative transporters (50); CD39/CD73 inhibitors would not suppress this type of *de novo* adenosine production.

With regard to the effect of adenosine on other Th subsets, our observations were different from those of previous reports (51, 52); here, we observed that adenosine upregulated IFN-γ (a Th1-related cytokine) secretion at 5 and 7 days and had no significant effect on IL-5 (a Th2-related cytokine) secretion by CD4^+^ T cells after CD3/CD28 stimulation with 600 μM of adenosine, although IL-17A production was significant (Sup. Fig. 3). This suggests that adenosine induces hypersecretion of IL-17A by Th17 cell but does not suppress Th1 and Th2 activity. However, previous studies report that adenosine-mediated suppression of IFN-γ and IL-5 was observed 1 day after T cell receptor-mediated stimulation of CD4^+^ T cells (51, 52). This may indicate that in the short term adenosine prioritizes stimulation of Th17 cell activity rather than that of Th1 and Th2 cells, and that it does not suppress effector Th activity in the long term.

It is also suggested that the A2aR agonist, CGS21680, suppresses Th17 differentiation (15–17). This result is opposite to ours; one reason for this may be differences in the source of the A2aR agonist. The A2aR agonist CGS 21680 is much less selective than the A2aR agonist that we used in this study (PSB0777); this is because CGS21680 binds not only to the A2aR but also to A1R and A3R, which are associated with the Gi protein (which has opposite effects to the Gs protein) (53, 54). Therefore, it is probable that CGS21680 may cancel out any agonist effects by activating A1R and A3R. Also, it is suggested that the A2aR antagonist, SCH58261, up-regulates Th17 differentiation in mice (17). We hypothesized that up-regulation of Th17 differentiation by SCH58261 may be induced through relative downregulation of A2aR activity compared with that of A2bR; this relative increase in A2bR activity induces Th17 differentiation in mice (55, 56). Also, it is probable that SCH58261 induces relative increase in the activity of G protein-coupled receptors other than adenosine receptors (e.g., dopamine receptors) to induce Th17 differentiation in mice (57). We also hypothesize that although SCH58261 may induce Th17 differentiation *in vivo*, it may not stimulate Th17 activity; this is because our data show that an A2a antagonist (Istradefylline) suppressed IL-17A secretion by differentiated Th17 cells but did not suppress Th17 differentiation (Fig. 5B). This hypothesis is supported by previous data showing that SCH58261 markedly suppresses symptoms of EAE, a typical Th17-mediated disease (34).

Our data suggest that production of IL-17A is relatively higher after exposure to an A2aR agonist, PSB0777, than after exposure to an A2bR agonist, BAY 60-6583. PSB0777 is a potent adenosine A2aR agonist (Ki = 44.4 nM for rat brain striatal A2aR) (54), and BAY 60-6583 is a potent adenosine A2bR agonist (Ki = 100 nM for rat A2bR) (58). By assuming that the Ki values of PSB0777 and BAY 60-6583 are comparable, we thought that production of IL-17A mediated by activation of the A2aR might be higher than that mediated by activation of the A2bR. This hypothesis is supported by the notion that the A2aR is a high affinity receptor with activity in the low to mid-nanomolar range, whereas the A2bR has a much lower affinity for adenosine (micromolar) (5); this suggests that adenosine activates the A2aR rather than the A2bR.

A2aR antagonists have been developed for treatment of Parkinsonism (59) and malignancies (60). In addition, inhibitors of CD39/CD73 have been developed as anti-tumor drugs (61, 62). Regarding the effects of adenosine on tumor immunity, a previous study suggests that adenosine suppresses effector T cell function since tumor cells express both CD39 and CD73 and secrete adenosine (63). By contrast, several reports suggest that IL-17A promotes emergence of pro-tumorigenic neutrophil phenotypes (64, 65). Neutrophils in mouse tumor models promote tumor metastasis (66–68), and observations in cancer patients have linked elevated neutrophil counts in blood with increased risk of metastasis (69). Therefore, it is probable that tumor-produced adenosine induces IL-17A secretion by CD4^+^CCR6^hi^ T cells followed by neutrophilic inflammation, which promotes tumor metastasis. Cancer vaccines may need to be administered along with an A2aR antagonist to suppress hypersecretion of IL-17A by tumor-specific Th17 cells induced by tumor-produced adenosine, and to inhibit neutrophilic inflammation at the tumor site.

The results presented herein indicate that these drugs may also be effective treatments for Th17-mediated diseases (4) such as psoriasis, neutrophilic bronchial asthma, severe atopic dermatitis, and autoimmune diseases by suppressing hypersecretion of IL-17A by Th17 cells. Moreover, these drugs might be effective treatments for diseases caused by neutrophilic inflammation of unknown cause in the dermis; such diseases include Behcet uveitis (70) and vasculitis of adenosine deaminase 2 deficiency (71). This is because endothelial cells express CD39/CD73 and produce adenosine, which could induce hypersecretion of IL-17A by Th17 cells, thereby contributing to inflammation.

## Abbreviations

Th: T-helper

TGF: tumor growth factor

IL: interleukin

APCs: antigen presenting cells

AC: adenyl cyclase

PKA: protein kinase A

CREB: cAMP response element binding protein

MLR: two-way mixed lymphocyte reaction

EAE: experimental autoimmune encephalomyelitis

CCPA: 2-Chloro-N6-cyclopentyladenosine

AMP-CP: adenosine 5’-(α, β-methylene) diphosphate

BM: bone marrow

BM-DC: bone-marrow-derived dendritic cell

CD3/CD28: agonistic anti-CD3/CD28 antibodies

PLP: myelin proteolipid protein

PLP peptide: I-A^s^ restricted helper peptide derived from the PLP

CFA: complete Freund’s adjuvant

CCR: chemokine receptor

FITC: fluorescein isothiocyanate

PE: phycoerythrin

MACS: magnetic-activated cell sorting

Ab: antibody

n: number of repeat experiments

SD: standard deviation.

## Acknowledgments

This work was supported by a Grant-in-Aid for Scientific Research (C) (no. 19K07201), awarded to M.K., a Grant-in-Aid for Young Scientists (B) (no. 18K15327) to R.T., and a Grant-in-Aid for Scientific Research (C) (no. 19K08887) awarded to S.M. by the Japanese Society for the Promotion of Science. This work was also supported by the 44th and 45th Science Research Promotion Fund, awarded to M.K. by the Promotion and Mutual Aid Corporation for Private Schools of Japan.

## Author contributions

M.T., R.T., S.M., and M.K., performed the experiments. M.T., S.M., and M.K., conceived and designed the experiments. M.T., S.M., T.Y., and M.K., wrote the manuscript. All authors discussed the results and commented on the manuscript.

## Legends for the supplementary figures

**Sup. Fig. 1.**
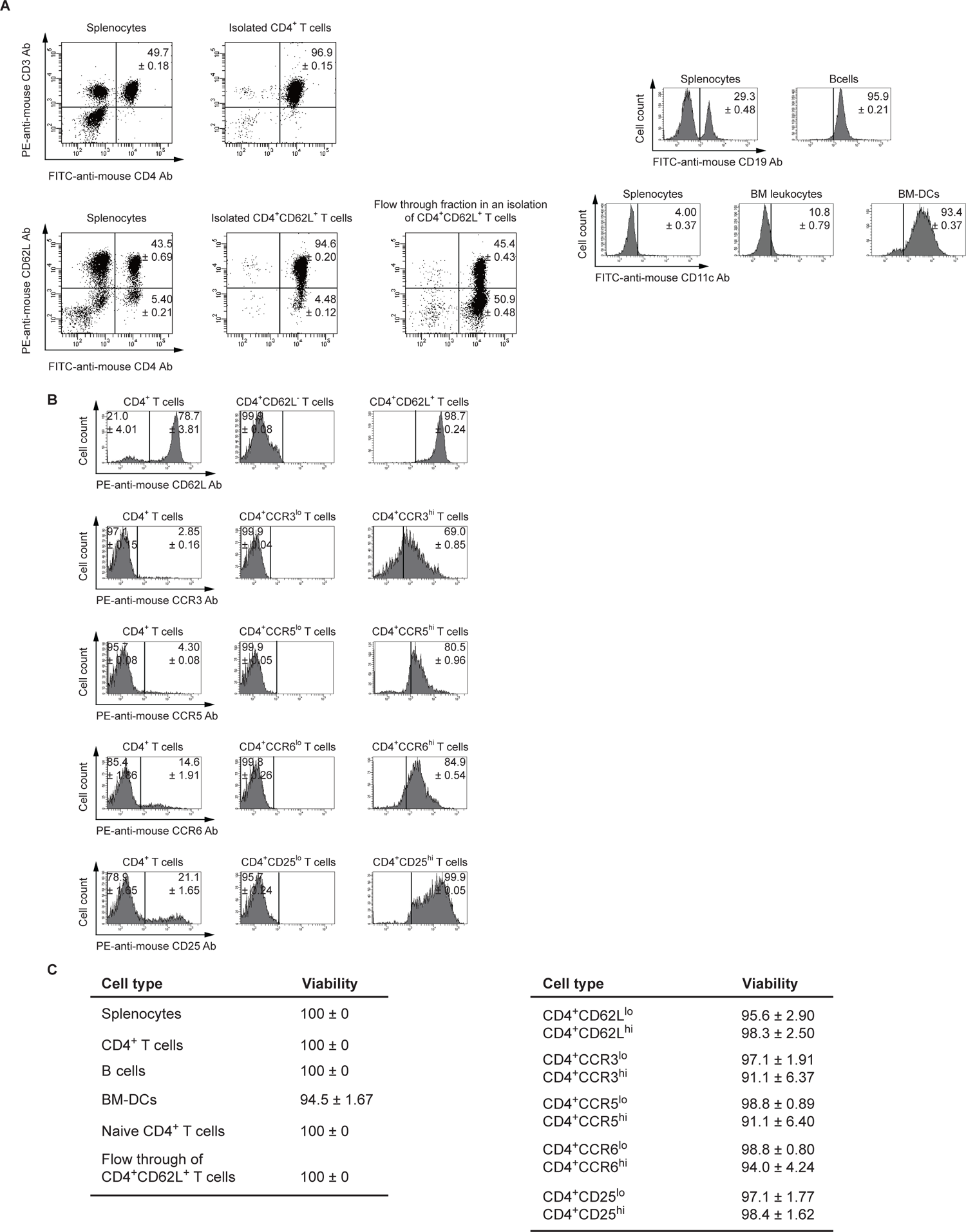
Purity and viability of each immune cell subset. **A**, *First row (left)*: Splenocytes or isolated CD4^+^ T cells were stained with a FITC-conjugated anti-mouse CD4 antibody (Ab) (BioLegend) (x-axis) and a PE-conjugated anti-mouse CD3 Ab (BioLegend) (y-axis). *Second row (left)*: Splenocytes or CD4^+^CD62L^+^ T cells isolated by MACS were stained with a FITC-conjugated anti-mouse CD4 Ab (BioLegend) (x-axis) and a PE-conjugated anti-mouse CD62L Ab (BioLegend) (y-axis). *First row (right)*: Splenocytes and B cells were stained with a FITC-conjugated anti-mouse CD19 Ab (BioLegend) (x-axis) and cell numbers were counted (y-axis). *Second row (right)*: Splenocytes, BM leukocytes, and BM-DCs were stained with a FITC-conjugated anti-mouse CD11c Ab (BioLegend) (x-axis) and cell numbers were counted (y-axis). The number in each panel represents the percentage ± SD of each immune subset within the total cell population (n = 3). **B**, *First row*: Isolated CD4^+^ (by MACS)-, CD4^+^CD62L^-^ (by cell sorting)-, or CD4^+^CD62L^+^ (by cell sorting) T cells were analyzed with a PE-conjugated anti-mouse CD62L Ab (BioLegend) (x-axis) and cell numbers were counted (y-axis). The number in each panel represents the percentage of each immune subset (- or +) within the total cell population (- plus +). Data are representative of at least three repeat experiments. *Second row*: Isolated CD4^+^ (by MACS), CD4^+^CCR3^low (lo)^ (by cell sorting), or CD4^+^CCR3^high (hi)^ (by cell sorting) T cells were analyzed with a PE-conjugated anti-mouse CCR3 Ab (BioLegend) (x-axis) and cell numbers were counted (y-axis). *Third row*: Isolated CD4^+^ (by MACS), CD4^+^CCR5^lo^ (by cell sorting), or CD4^+^CCR5^hi^ (by cell sorting) T cells were analyzed with a PE-conjugated anti-mouse CCR5 Ab (BioLegend) (x-axis) and cell numbers were counted (y-axis). *Fourth row*: Isolated CD4^+^ (by MACS), CD4^+^CCR6^lo^ (by cell sorting), or CD4^+^CCR6^hi^ (by cell sorting) T cells were analyzed with a PE-conjugated anti-mouse CCR6 Ab (BioLegend) (x-axis) and cell numbers were counted (y-axis). *Fifth row*: Isolated CD4^+^ (by MACS), CD4^+^CD25^lo^ (by cell sorting), or CD4^+^CD25^hi^ (by cell sorting) T cells were analyzed with a PE-conjugated anti-mouse CD25 Ab (BioLegend) (x-axis) and cell numbers were counted (y-axis). The number in each panel represents the mean percentage ± SD of each immune subset (hi or lo) within the total cell population (hi plus lo) (n = 3). **C**, After cell isolation, each immune cell type was mixed with Trypan blue. Viability was calculated as the number of unstained cells/(stained cells + unstained cells) × 100 (n = 3). The percentage represents the mean percentage ± SD.

**Sup. Fig. 2.**
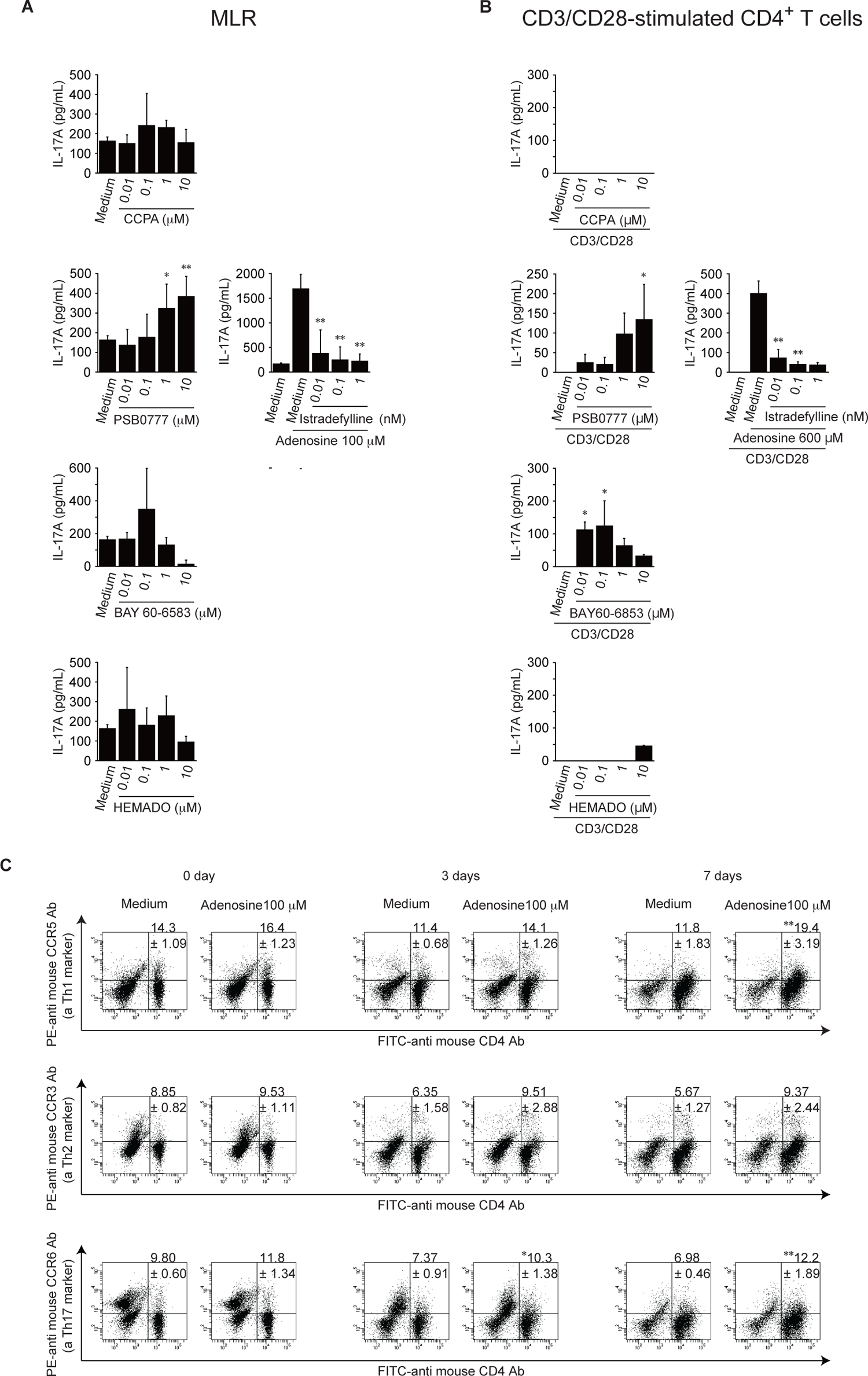
Effect of an adenosine receptor agonist and antagonist on production of IL-17A in an MLR and by CD3/CD28-stimulated CD4^+^ T cells, and the effect of adenosine on proliferation of Th cells. A, An MLR was performed for 7 days in the presence of each adenosine receptor agonist, or each adenosine receptor antagonist plus adenosine (100 μM) (n = 6–9). B, CD4^+^ T cells were stimulated with an anti-CD3/CD28 antibody for 7 days in the presence of each adenosine receptor agonist, or in the presence of each adenosine receptor antagonist plus adenosine (600 μM) (n = 6–9). After 7 days, the supernatants were analyzed in an IL-17A ELISA. *First row*: CCPA (an A1R agonist). *Second row*: PSB0777 (an A2aR agonist, *left)* and Istradefylline (an A2aR antagonist, *right*). *Third row*: BAY 60-653 (an A2bR agonist). *Forth row*: HEMADO (an A3R agonist). C, An MLR was performed in the presence or absence of adenosine (100 μM). After 0, 3, and 7 days, splenocytes were stained with anti-CD4 (x-axis) and CCR5 (a Th1 marker; top panels, y-axis), CCR3 (a Th2 marker; second panels, y-axis), or CCR6 (a Th17 marker; bottom panels, y-axis) antibodies, followed by flow cytometry analysis (n = 3). Values in the upper right of each panel denote the percentage of CCR-positive counts within the CD4 positive counts. *P < 0.05 and **P < 0.01. Data are expressed as the mean ± SD and are compared using one-way ANOVA with Tukey’s post-hoc test. *P < 0.05 and **P < 0.01, compared with medium (A, *left*; B, *left*; and C), adenosine (100 μM) (A, *right*), CD3/28 stimulation plus adenosine (600 μM) (B, *right*).

**Sup. Fig. 3.**
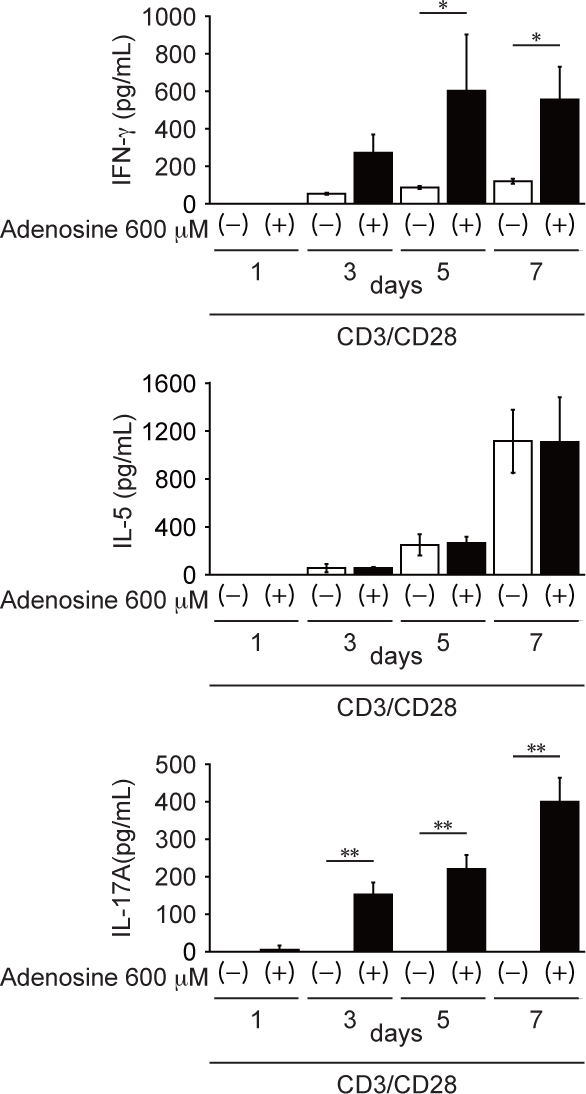
Effect of adenosine on production of IFN-γ, IL-5, and IL-17A by CD3/CD28-stimulated CD4^+^ T cells. CD4^+^ T cells were stimulated for 1–7 days with anti-CD3/CD28 antibodies in the presence or absence of adenosine (600 μM). After stimulation, supernatants were analyzed in IFN-γ (*top row*), IL-5 (*second row*), and IL-17A (*bottom row*) ELISAs (n = 4–6). Data are expressed as the mean ± SD and were compared using one-way ANOVA with Tukey’s post-hoc test. *P < 0.05 and **P < 0.01, compared with CD3/CD28 stimulation.

**Sup. Fig. 4.**
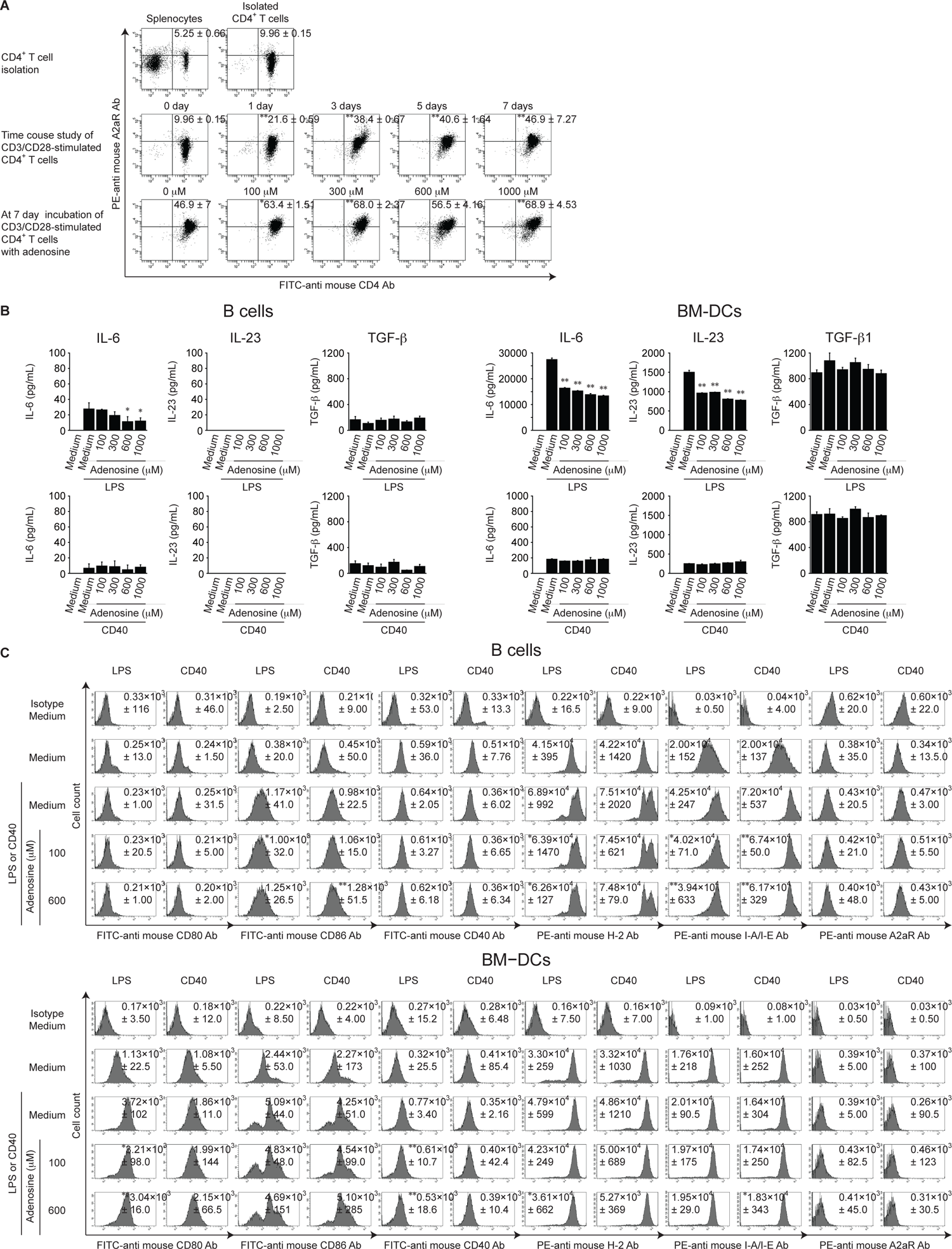
Effect of adenosine on A2aR expression by CD4^+^ T cells, and on expression of MHC, costimulatory molecules, and A2aR and production of IL-6, IL-23, and TGF-β by APCs. A, CD4^+^ T cells were stimulated for 0–7 days with anti-CD3/CD28 antibodies in the presence of adenosine (0–1 mM). After stimulation, cells were stained with a FITC-conjugated anti-mouse CD4 antibody (Ab) (BioLegend) (x-axis) and a PE-conjugated anti-A2aR Ab (Novus Biologicals) (y-axis). *First row*: Splenocytes (*left*) or isolated CD4^+^ T cells (*right*) were stained with anti-CD4 (x-axis) and A2aR (y-axis) Abs. *Second row*: CD3/CD28-stimulated CD4^+^ T cells were stained with anti-CD4 (x-axis) and A2aR (y-axis) Abs. *Third row*: CD4^+^ T cells were stimulated with CD3/CD28 Abs in the presence of adenosine (0–1 mM) and then stained with anti-CD4 (x-axis) and A2aR (y-axis) Abs. The number in each panel represents the percentage ± SD of A2aR^high^ cells within the CD4^+^ population (n = 3). Data were compared using one-way ANOVA with Tukey’s post-hoc test. *P < 0.05 and **P < 0.01, compared with CD3/CD28 stimulation (second and third rows). B, B cells and BM-DCs were stimulated for 1 day with LPS (upper panels) or anti-mouse CD40 Ab (CD40) (lower panels) in the presence of adenosine (0.1– 1 mM). After stimulation, the IL-6, IL-23, and TGF-β1 concentrations in the supernatants were analyzed by ELISA (n = 3). Data are expressed as the mean ± SD and were compared using one-way ANOVA with Tukey’s post-hoc test. *P < 0.05 and **P < 0.01, compared with LPS or anti-mouse CD40 Ab stimulation. C, B cells (upper panels) and BM-DCs (lower panels) were stimulated for 1 day with LPS or anti-mouse CD40 Ab (CD40) in the presence of adenosine (100 or 600 μM). After stimulation, the cells were stained with a FITC-conjugated anti-mouse CD80 (BioLegend), CD86 (BioLegend), or CD40 (BioLegend) Ab, PE-conjugated anti-mouse H-2 (BioLegend) or I-A/I-E (BioLegend) Ab, or a PE-conjugated anti-A2aR (Novus Biologicals) Ab (x-axis), and cell numbers were counted (y-axis) (n = 3). The number in each panel represents the mean fluorescence intensity ± SD (n = 3). Data were compared using one-way ANOVA with Tukey’s post-hoc test. *P < 0.05 and **P < 0.01, compared with LPS or anti-mouse CD40 Ab stimulation.

